# Mechanical and molecular parameters that influence the tendon differentiation potential of C3H10T1/2 cells in 2D- and 3D-culture systems

**DOI:** 10.1101/760751

**Authors:** Ludovic Gaut, Marie-Ange Bonnin, Isabelle Cacciapuoti, Monika Orpel, Mathias Mericskay, Delphine Duprez

**Affiliations:** Sorbonne Université, Institut Biologie Paris Seine, CNRS, IBPS-UMR7622, Laboratoire de Biologie du Développement, Inserm U1156, F75005 Paris, France; Inovarion, Paris France; Inserm UMR-S 1180, Faculté de Pharmacie, Univ. Paris-SUD, Université Paris-Saclay, F-92296 Châtenay-Malabry, France

**Keywords:** Mesenchymal stem cells, Cell confluence, 2D- and 3D-cultures, Plastic substrate, Silicone substrate, Tendon differentiation, TGFβ2, Scleraxis, Tenomodulin

## Abstract

One of the main challenges in tendon field relies in the understanding of regulators of the tendon differentiation program. The optimum culture conditions that favor tendon cell differentiation are not identified. Mesenchymal stem cells present the ability to differentiate into multiple lineages in cultures under different cues ranging from chemical treatment to physical constraints. We analyzed the tendon differentiation potential of C3H10T1/2 cells, a murine cell line of mesenchymal stem cells, upon different 2D- and 3D-culture conditions. We observed that C3H10T1/2 cells cultured in 2D conditions on silicone substrate were more prone to tendon differentiation assessed with the expression of the tendon markers *Scx, Col1a1* and *Tnmd* as compared to cells cultured on plastic substrate. 3D fibrin environment was more favorable for *Scx* and *Col1a1* expression compared to 2D-cultures. We also identified TGFβ2 as a negative regulator of *Tnmd* expression in C3H10T1/2 cells in 2D- and 3D-cultures. Altogether, our results provide us with a better understanding of the culture conditions that promote tendon gene expression and identify mechanical and molecular parameters on which we could play to define the optimum culture conditions that favor tenogenic differentiation in mesenchymal stem cells.

## Introduction

Mesenchymal stem cells (MSCs) are multipotent cells that can be induced to differentiate in various tissue lineages upon specific molecular or mechanical cues. Based on specific lineage markers and identified master genes, established protocols are now recognized to drive differentiation towards osteocytes, chondrocytes and adipocytes (Caplan, 1991; Pittenger et al., 1999; Prockop, 1997). Although studies identify tendon cell differentiation upon molecular and mechanical cues from MSCs (reviewed in Nourissat et al., 2015; Zhang et al., 2018), the tendon lineage is understudied compared to other tissue-specific lineages. There is no recognized/established protocol with external inducers to differentiate MSCs towards a tendon phenotype. In addition, there is no identified master gene that initiates the tenogenic program in cell cultures as for the cartilage (Sox9), bone (Runx2) and muscle (Muscle regulatory factors) programs (Buckingham, 2017; Karsenty et al., 2009; Liu et al., 2017).

Another difficulty to study tendon differentiation is the limited number of specific tendon markers. The main structural and functional component of tendon, type I collagen is not specific to tendon and is expressed in many other connective tissues (reviewed in Gaut and Duprez, 2016). To date, the bHLH transcription factor Scleraxis (Scx) is the best marker for tendons and ligaments during development (Schweitzer et al., 2001; Schweitzer et al., 2010) and in the adult (Mendias et al., 2012). Although being a powerful tendon marker, the exact function of *Scx* in tendon development, homeostasis and repair is still not fully understood (Huang et al., 2015; Murchison et al., 2007). The type II transmembrane glycoprotein tenomodulin, encoded by the *Tnmd* gene, is recognized to be a tendon differentiation marker with potential roles in tenocyte proliferation and differentiation, in addition to type I collagen fibril adaptation to mechanical loads (Alberton et al., 2015; Dex et al., 2016; Dex et al., 2017; Docheva et al., 2005). *Scx* is required for *Tnmd* expression in mouse tendons during development (Murchison et al., 2007; Yoshimoto et al., 2017). Scx gain- and loss-of-function experiments combined with electrophoresis mobility shift assay (EMSA) in cell cultures indicate a direct regulation of Scx on *Tnmd* promoter (Shukunami et al., 2018; Yoshimoto et al., 2017). In addition to the well-studied tendon markers, *Scx* and *Tnmd*, a list of 100 tendon markers has been identified in limb tendon cells during mouse development via transcriptomic analysis (Havis et al., 2014).

The main extracellular signal known to promote tendon development is the TGFβ ligand (Havis et al., 2014, 2016; Maeda et al., 2011; Pryce et al., 2009). TGFβ ligands are recognized to have a generic protenogenic effect based on the increase of *Scx* transcription in cell cultures (Guerquin et al., 2013; Havis et al., 2014; Havis et al., 2016; Lorda-Diez et al., 2009; Pryce et al., 2009). The increase of *Scx* expression upon TGFβ2 exposure is abolished in the presence of TGFβ inhibitors, which block TGFβ signal transduction at the level of the receptors or at the level of the SMAD2/3 intracellular pathways in C3H10T1/2 cells (Guerquin et al., 2013; Havis et al., 2014).

In addition to chemical signals, mechanical signals are important parameters to consider when studying tendon cell differentiation. Because tendons transmit forces from muscle to bone in the musculoskeletal system, tendon cells are continuously subjected to variations in their mechanical environment (Schiele et al., 2013). Physical constraints subjected to the cells have been shown to be important for developmental processes and during adult life (Mammoto et al., 2013). It is recognized that substrate stiffness controls many cellular processes such as cell fate, migration, proliferation and differentiation in culture systems of stem cells or progenitor cells (Bellas and Chen, 2014; Ivanovska et al., 2015; Kilian et al., 2010). MSCs are particularly responsive to matrix stiffness in term of lineage commitment, ranging from neurogenic phenotype for soft substrates to osteogenic when cultured on rigid substrates (Discher et al., 2009; Engler et al., 2006; Humphrey et al., 2014). The forces transmitted through cell contacts upon confluence is another parameter that mechanically constrains cells in culture dishes and influences cell differentiation (Abo-Aziza and Zaki, 2017; Ren et al., 2015).

The tendon phenotype is not maintained in 2D-cultures of tendon cells over passages (Hsieh et al., 2018; Shukunami et al., 2018; Yao et al., 2006). 3D-culture systems, in which tendon cells are embedded in hydrogels are recognized to provide an environment closer to that experienced by tendon cells *in vivo* (Kapacee et al., 2010; Kuo et al., 2010; Marturano et al., 2016; Yeung et al., 2015). The mechanical environment provided to tendon cells homogeneously embedded within hydrogel in 3D-culture systems is recognized to act on tendon gene expression (Hsieh et al., 2018; Marturano et al., 2016). Most of the analyses of the effects of 2D- and 3D- environments have been performed with tendon stem/progenitor cells; however, the optimum culture conditions that drive tendon cell differentiation from MSCs have not been yet identified. In the present study, we analyzed the tendon differentiation potential of C3H10T1/2 cells under different mechanical and molecular signals in 2D- and 3D- culture conditions.

## Results

In order to investigate the tendon differentiation potential, we used C3H10T1/2 cells, a multipotent cell line established from mouse embryos (Reznikoff et al., 1973). C3H10T1/2 cells are known to differentiate into chondrocytes, osteocytes and adipocytes when cultured under appropriate cues (Guerquin et al., 2013). These cells have the ability to display a tendon phenotype under inductive molecular cues, such as the transcription factors EGR1 and MKX (Guerquin et al., 2013; Liu et al., 2015). The ability to differentiate into cell lineages related to the musculoskeletal system makes of the C3H10T1/2 cells an ideal tool to study tendon commitment and differentiation under different mechanical and molecular cues in 2D- and 3D- culture conditions. To assess tendon differentiation, we used the mRNA levels of key tendon markers, *Scx* and *Tnmd* in addition to *Col1a1,* the main structural and functional tendon component. We also used tendon genes identified in the transcriptomic analysis of mouse tendon cells during development (Havis et al., 2014), such as aquaporin1 (*Aqp1*) gene coding for a water channel protein and thrombospondin 2 (*Thsb2)* coding for an adhesive glycoprotein with antiangiogenic properties, both expressed in developing limb tendons.

### Seeding density does not affect tendon gene expression in non-confluent conditions after 16 hours of culture

We first determined whether the initial cell number interfered with the expression of tendon genes in non-confluent conditions. Different cell numbers, 0.5×10^5^, 10^5^ and 2×10^5^ cells, were seeded in 9 cm^2^ culture plates (plastic substrate), corresponding to 5 555 cells/cm^2^, 11 110 cells/cm^2^ and 22 220 cells/cm^2^, respectively. After 16 hours of culture, the expression of tendon genes, *Scx*, *Tnmd*, *Col1a1* and *Aqp1* did not display any change more than 20% upon different cell density seeding conditions (Figure 1AB). This shows that the initial cell number at seeding time does not have a major influence on tendon gene expression in expansion and non-confluent conditions.

**Figure 1.**
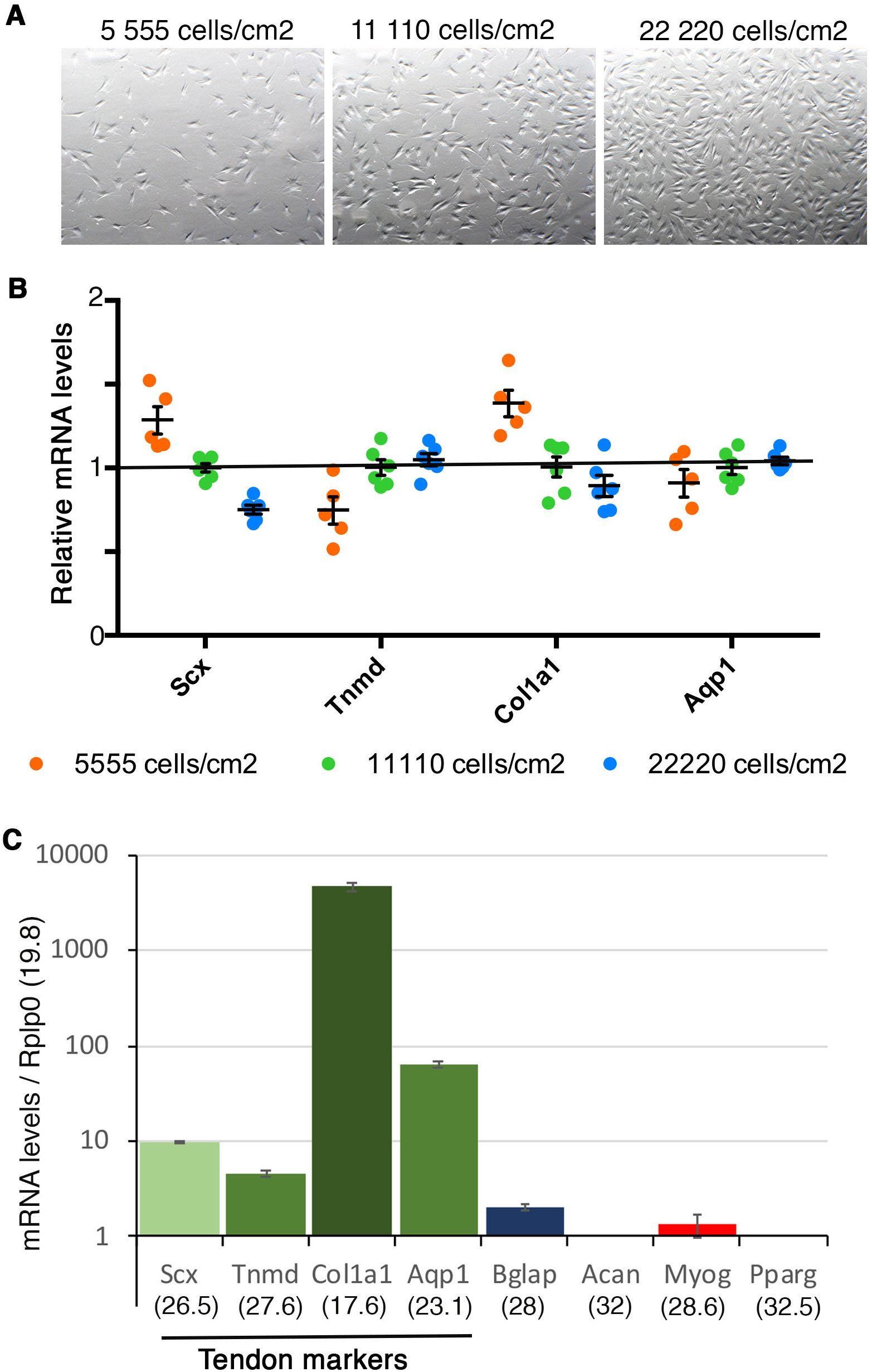
Tendon gene expression is not related to cell density in non-confluent conditions. **(A)** Representative pictures of cell density 16 hours after plating 0.5×10^5^ (5 555 cells/cm^2^), 10^5^ (11 111 cells/cm2) and 2×10^5^ (22 222 cells/cm^2^) C3H10T1/2 cells on 9 cm^2^ plastic culture plates. **(B)** RT-q-PCR analyses of the expression levels of tendon markers, *Scx, Tnmd, Col1a1* and *Aqp1* in C3H101/2 cells 16 hours after initial plating of 5 555 cells/cm^2^, 11 110 cells/cm^2^ and 22 220 cells/cm^2^. The relative mRNA levels were calculated using the 2^^-ΔΔCt^ method using the 11 110 cells/cm^2^ plating condition as controls. For each gene, the mRNA levels of the 11 110 cells/cm^2^ plating condition were normalized to 1 (green spots). Graph shows means ± standard deviations of 6 biological samples. **(C)** RT-q-PCR analyses of the expression levels for the tendon markers, *Scx, Tnmd, Col1a1*, *Aqp1* and for lineage markers, *Bglap* (bone), *Acan* (cartilage), *Myog* (muscle) and *Pparg* (fat) in C3H101/2 cells 16 hours after initial plating 11 110 cells/cm^2^ on plastic culture plates. mRNA levels on the Y-axis are reported to the *Rplp0 (36b4)* gene (2^^-ΔCt^ x 10^3^). Graph shows means ± standard deviations of 6 biological samples. The means of the initial Cts (obtained from 250 ng of mRNAs) are indicated in brackets for each gene. *Rplp0* (Cts=19.6 SD+/-0.17); *Col1a1* (Cts=17.6, SD+/- 0.2); *Aqp1* (Cts= 23.1 SD+/-0.44), *Scx* (Cts=26.5 SD+/-0.22; *Tnmd* (Cts=27.6 SD+/-0.28); *Bglap* (bone, Cts=28 +/-0.38); *Myog* (muscle, Cts=28.6+/-0.16). *Acan* (cartilage) and *Pparg* (fat) displayed Cts above 32.

We next compared the relative mRNA expression levels between tendon genes in C3H10T1/2 cells in non-confluent conditions on plastic substrate (at the 11 110 cells/cm^2^ seeding condition). The expression levels of each tendon gene were reported to the *Rplp0* gene (Ct= 19.8 for 250 ng of RNAs). We found that *Col1a1* gene display high expression levels (Ct=17.6) compared to those of *Aqp1* (Ct= 23.1), *Scx* (Ct=26.5) and *Tnmd* (Ct=27.6) genes in C3H10T1/2 cells in non-confluent condition. Analysis of the mRNA expression levels for other lineage markers showed that *Acan* (cartilage) and *Pparg* (fat) genes were not expressed (Ct above 32), while *Bglap* (bone, Ct=28) and *Myog* (muscle, Ct=28.6) genes displayed low levels of expression in C3H10T1/2 cells in non-confluent conditions on plastic substrate (Figure 1C). This shows that tendon genes are expressed in C3H10T1/2 cells seeded in non-confluent conditions on plastic substrate after 16 hours of culture, with an expression level superior to that of other differentiation markers such as bone, cartilage, muscle and fat. We conclude that C3H10T1/2 cells display a fibroblastic phenotype.

### Gene expression profiles are similar in C3H10T1/2 cells seeded on two different substrates on the rigid scale in non-confluent conditions after 16 hours of culture

The same density of C3H10T1/2 cells (11 110 cells/cm^2^) was plated on classic culture plastic plates and Uniflex Flexcell plates made of silicone substrate coated with type I collagen (Figure 2A,B). Plastic substrate displays a Young Modulus of 1 GPa magnitude and is considered as extremely rigid. Uniflex Flexcell plates display a stiffness estimated of 5 MPa by the company (Flexcell international corporation). The silicon substrate is 200-fold less rigid (5 MPa) compared to plastic substrate (1 GPa) but is still considered as rigid on the micro-stiffness scale for substrates (Discher et al., 2009). C3H10T1/2 cells were harvested 16 hours after plating at a non-confluent state (Figure 2A,B) and similar amount of mRNAs was analyzed for gene expression. Tendon and other lineage marker expression profiles were similar in both substrate culture conditions (Figure 2C,D). This shows that two substrates with different stiffnesses on the rigid scale do not affect gene expression profiles in C3H10T1/2 cells seeded in non-confluent conditions for 16 hours.

**Figure 2.**
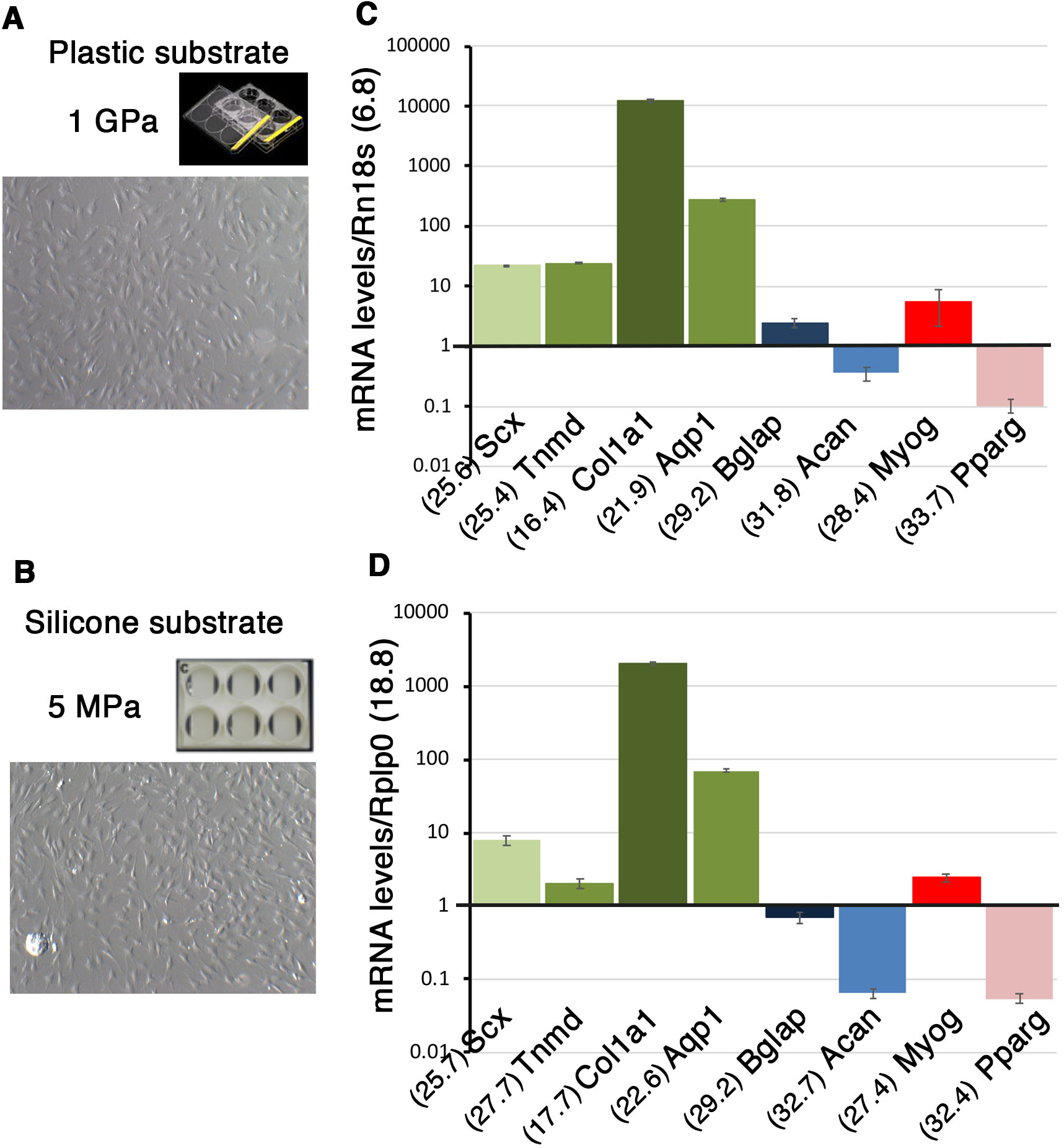
The nature of the substrate does not modify gene expression profiles in C3H10T1/2 cells in non-confluent conditions. (**A,B**) Photographs of C3H10T1/2 cells cultured on plastic plates displaying a stiffness of 1 GPa (A) and on Uniflex culture plates made of silicon coated with type I collagen displaying a stiffness of 5 Mpa (B), in non-confluent conditions. (**C,D**) RT-q-PCR analyses of gene expression levels in C3H10T1/2 cells. *Scx, Tnmd, Col1a1*, *Aqp1* and representative genes for the bone (*Bglap*), cartilage (*Acan*), muscle, (*Myog)* and fat (*Pparg)* lineages, in C3H101/2 cells in non-confluent conditions on plastic (C) and silicone (D) substrates. The means of the Cts (obtained from 500 ng of mRNAs) are indicated in brackets for each gene. (C) Plastic substrate: for each gene, the ΔCt was calculated using *Rn18S* as reference gene. ΔCt= Ct *gene* - Ct *Rn18S*. The mRNA levels were reported using the 2^^-ΔCt^ method. In order to obtain values above 1, each 2^^-ΔCt^ were multiplied per 10^6^. Graph shows means ± standard deviations of 4 biological samples. (D) Silicone substrate: for each gene, the ΔCt was calculated using the *Rplp0* gene as reference gene. ΔCt = Ct *gene* - Ct *Rplp0.* The mRNA levels were calculated using the 2^^-ΔCt^ method. For each gene, 2^^-ΔCt^ were multiplied per 10^3^. Graph shows means ± standard deviations of 6 biological samples.

### Differentiation potential of C3H10T1/2 cells cultured on plastic substrate over time

We investigated the tendon differentiation potential of C3H10T1/2 cells cultured on plastic substrate over time. Cells were plated on plastic culture plates at 11 110 cells/cm^2^ density and left for 16 hours to define the Day 0. C3H10T1/2 cells were let to grow for 14 days with no passage. C3H10T1/2 cells were harvested at 1 day, 7 days, 10 days and 14 days of culture. The cell density of C3H10T1/2 cells was measured (Figure 3A,B) at each time point. At Day 0 (16 hours after plating) we obtained 17 100 cells/cm^2^ (SD +/- 4885, N=12). Cells expanded until Day 10 and reached a plateau from Day 10 to Day 14, defining 2 phases, one expansion phase until Day 10 and a post-expansion phase after Day 10 (Figure 3B).

**Figure 3.**
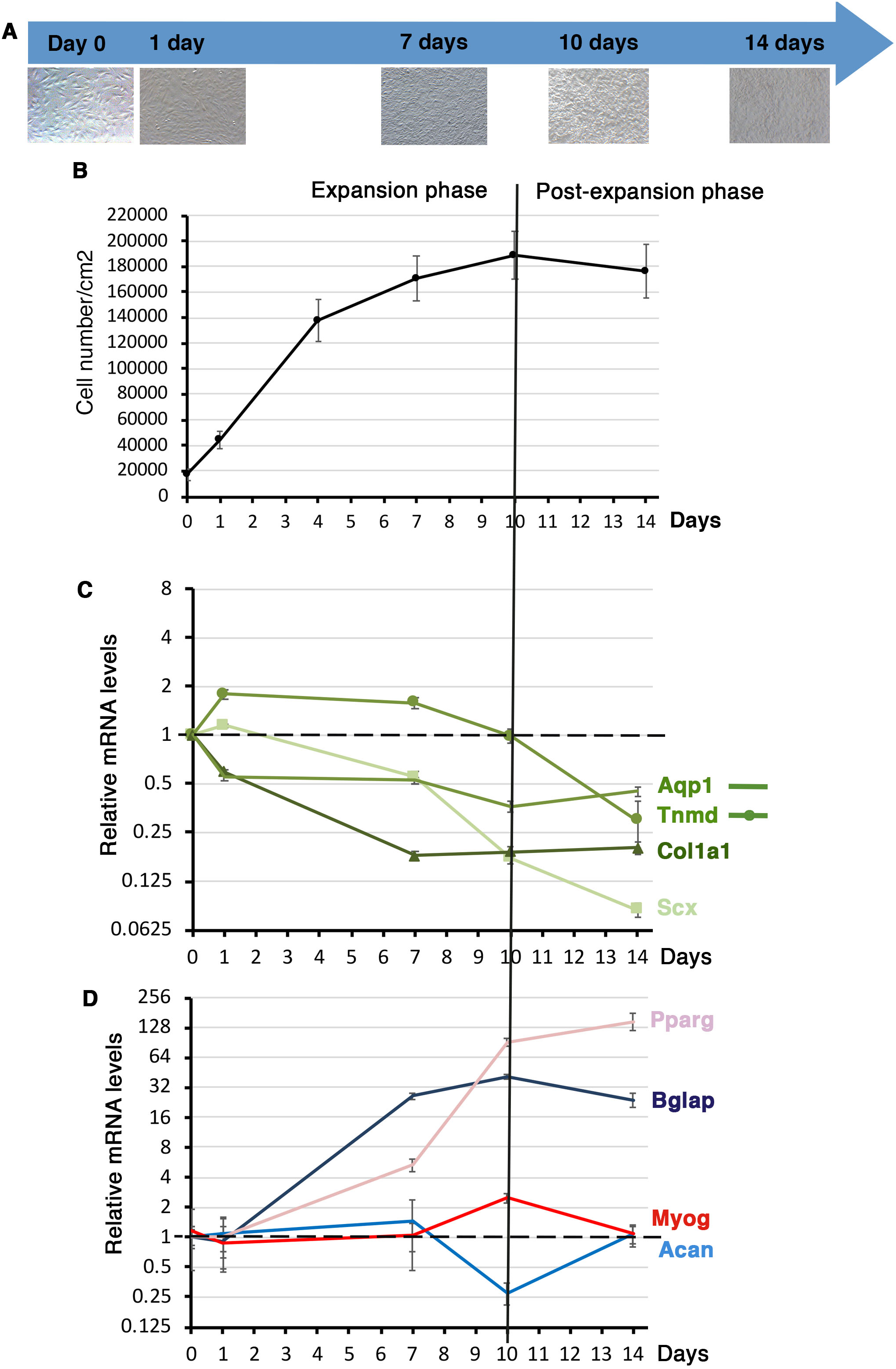
Gene expression in C3H10T1/2 cells cultured on plastic substrate over time. **(A)** Photographs of C3H101/2 cells cultured on plastic culture plates at different time points. 10^5^ C3H101/2 cells were plated on plastic culture plates and left for 16 hours to define the T=0 time point. Cells were then fixed at 1 day, 7 days, 10 days and 14 days for RT-q-PCR analyses. (**B**) Cell density or cell number/cm2 was measured at each time point. (**C,D**) RT-q-PCR analyses of the expression levels for tendon genes, *Scx, Tnmd, Col1a1*, *Aqp1* (C) and other cell lineage markers, *Bglap* (bone), *Acan* (cartilage), *Myog* (muscle) and *Pparg* (fat) in C3HT101/2 cells cultured on plastic culture plates at different time points. Gene mRNA levels were normalized to *Rplp0*. The relative mRNA levels were calculated using the 2^^-ΔΔCt^ method using the Day 0 condition as controls. For each gene, the mRNA levels of the T=0 condition were normalized to 1. Graph shows means ± standard deviations of 4 biological samples for T=0, 1 day, 7 days and 14 days and of 5 biological samples for 10 days.

Lineage-specific gene expression analysis was conducted in order to assess the differentiation behavior of C3H10T1/2 cells cultured on plastic substrate over time. During the expansion phase (before Day 10), *Scx, Col1a1* and *Aqp1* genes displayed a continuous decrease of mRNA levels, while *Tnmd* mRNA levels displayed a bell shape with a maximum of 2-fold increase between Day 1 and Day 7 (Figure 3C). During the post-expansion phase, *Scx* and *Tnmd* expression decreased, while that of *Col1a1* and *Aqp1* was stable (Figure 3C). We also analyzed the expression of differentiation markers for other components of the musculoskeletal system, ranging from high to soft intrinsic tissue stiffness: bone (*Bglap, Pparg*), cartilage (*Acan*), muscle (*Myog*) and fat (*Pparg*) (Figure 3D). *Acan* and *Myog* genes did not show any massive changes of expression over time (Figure 3D), while the bone differentiation marker *Bglap* and the early fat differentiation marker *Pparg* displayed a striking increase of expression levels during the time of the culture (Figure 3D). It has to be noted that these results were reported to the Day 0 time point, when *Bglap* and *Pparg* were hardly expressed (Figure 2C). These results showed that confluence increased the expression of bone and fat differentiation markers, in C3H10T1/2 cells cultured on plastic substrate over time.

We conclude that cell expansion to confluence has a global negative effect on the expression of *Scx, Col1a1* and *Aqp1* tendon lineage markers, while promoting that of bone and fat markers in C3H10T1/2 cells cultured on plastic substrate.

### Differentiation potential of C3H10T1/2 cells culture on silicon substrate over time

We next investigated the tendon differentiation potential of C3H10T1/2 cells cultured on silicon substrate over time. Similarly to cultures on plastic substrate, 10^5^ cells were plated on the Uniflex Flexcell plates and left for 16 hours, which defined the Day 0. Cells were then cultured for 1 day, 2 days, 4 days, 7 days and 11 days with no passage. The cell density of C3H10T1/2 cells was measured at each time point (Figure 4A,B). At Day 0 we obtained 19 298 cells/cm^2^ (SD +/- 8 568, N=12) (Figure 4B). C3H10T1/2 cells expanded until 7 days of culture on silicone substrate and then stopped growing to reach a plateau, defining the expansion and post-expansion phases (Figure 4B).

**Figure 4.**
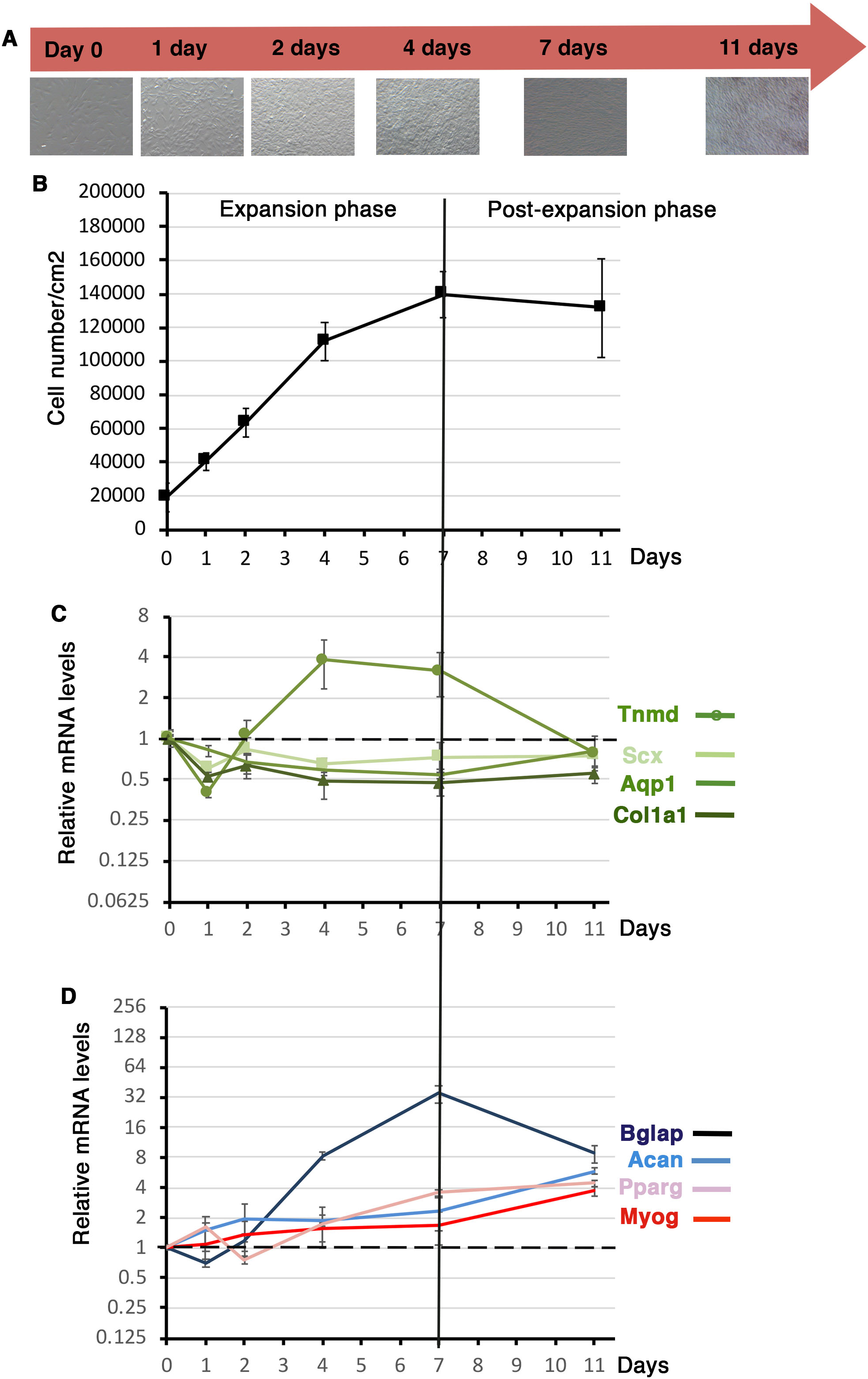
Gene expression in C3H10T1/2 cells cultured on silicone substrate over time. **(A)** C3H101/2 cells cultured on silicone substrate at different time points. 10^5^ C3H101/2 cells were plated on Uniflex Flexcell plates made of silicone and coated with type I collagen, and left for 16 hours to define the Day 0 time point. Cells were then fixed at 1 day, 2 days, 4 days, 7 days and 11 days for RT-q-PCR analyses. (**B**) Cell density or cell number/cm^2^ was measured at each time point. (**C,D**) RT-q-PCR analyses of the expression levels of tendon markers, *Scx, Tnmd, Col1a1*, *Aqp1* (**C**) and other cell lineage markers, *Bglap* (bone), *Acan* (cartilage), *Myog* (muscle) and *Pparg* (fat) (**D**) in C3HT101/2 cells cultured on silicone substrate at different time points. Gene mRNA levels were normalized to *Rplp0*. The relative mRNA levels were calculated using the 2^^-ΔΔCt^ method using the T=0 condition as controls. For each gene, the mRNA levels of the T=0 condition were normalized to 1. Graph shows means ± standard deviations of 5 biological samples for Day 0 and of 6 biological samples for 1 day, 2 days, 4 days, 7 days and 11 days.

Tendon gene expression analysis of C3H10T1/2 cells cultured on silicon substrate showed that the relative expression levels of all analyzed tendon genes, *Scx, Col1a1*, *Tnmd* and *Aqp1* decreased (up to 2-fold) during the first day of culture compared to Day 0 that was arbitrary normalized at 1 (Figure 4C). The *Scx*, *Col1a1* and *Aqp1* expression levels remained stable during of the rest of the culture but below the Day 0 expression levels (Figure 4C). The relative mRNA levels of the differentiation tendon gene *Tnmd* increased again after Day 1 and displayed a bell shape with a maximum of 4-fold increase between 4 and 7 days during the expansion phase and a decrease during the post-expansion phase (7 days to 11 days) (Figure 4C). This showed that the growing phase of C3H10T1/2 cells on silicone substrate was beneficial for *Tnmd* expression. As for plastic substrate, the bone marker *Bglap* displayed an increase of relative mRNA levels in C3H10T1/2 cells cultured on silicone substrate during the expansion phase (35-fold increase at 7 days relative to T=0) and decreased during the post-expansion phase (Figure 4D). The expression of the representative markers of differentiation for cartilage (*Acan*), muscle (*Myog*) and fat (*Pparg*) displayed a progressive increase over time to reach 4-fold at 11 days of culture (Figure 4D).

We conclude that the expansion phase has a positive effect on Tnmd gene expression in C3H10T1/2 cells cultured on silicone substrate over time.

### Tendon differentiation potential of C3H10T1/2 cells in a 3D-culture system

We next investigated the differentiation potential of C3H10T1/2 cells in a 3D-culture system. We used the 3D-fibrin gel method to produce in vitro-engineered tendons (Gaut et al., 2016; Guerquin et al., 2013; Kapacee et al., 2010). This 3D-culture system is based on tension (Bayer et al., 2010) and has been extensively characterized for matrix production by tendon progenitor cells (Yeung et al., 2015). We engineered 3D-fibrin constructs with C3H10T1/2 cells (Figure 5A-C). 3D-fibrin constructs took 5 to 7 days to fully form depending on the cultures (Figure 5A). The Day 0 was defined when the constructs was formed (Figure 5A,B). Transverse sections to a 24-hours construct show a homogeneous cell organization within the constructs (Figure 5C). We compared tendon gene expression in C3H10T1/2 cells cultured in 3D environment versus 2D plastic condition. The relative mRNA levels of *Scx* and *Col1a1* were significantly increased in C3H10T1/2 cells cultured in 3D versus 2D conditions, while those of *Tnmd* were not after 10 days of cultures (Figure 5D).

**Figure 5.**
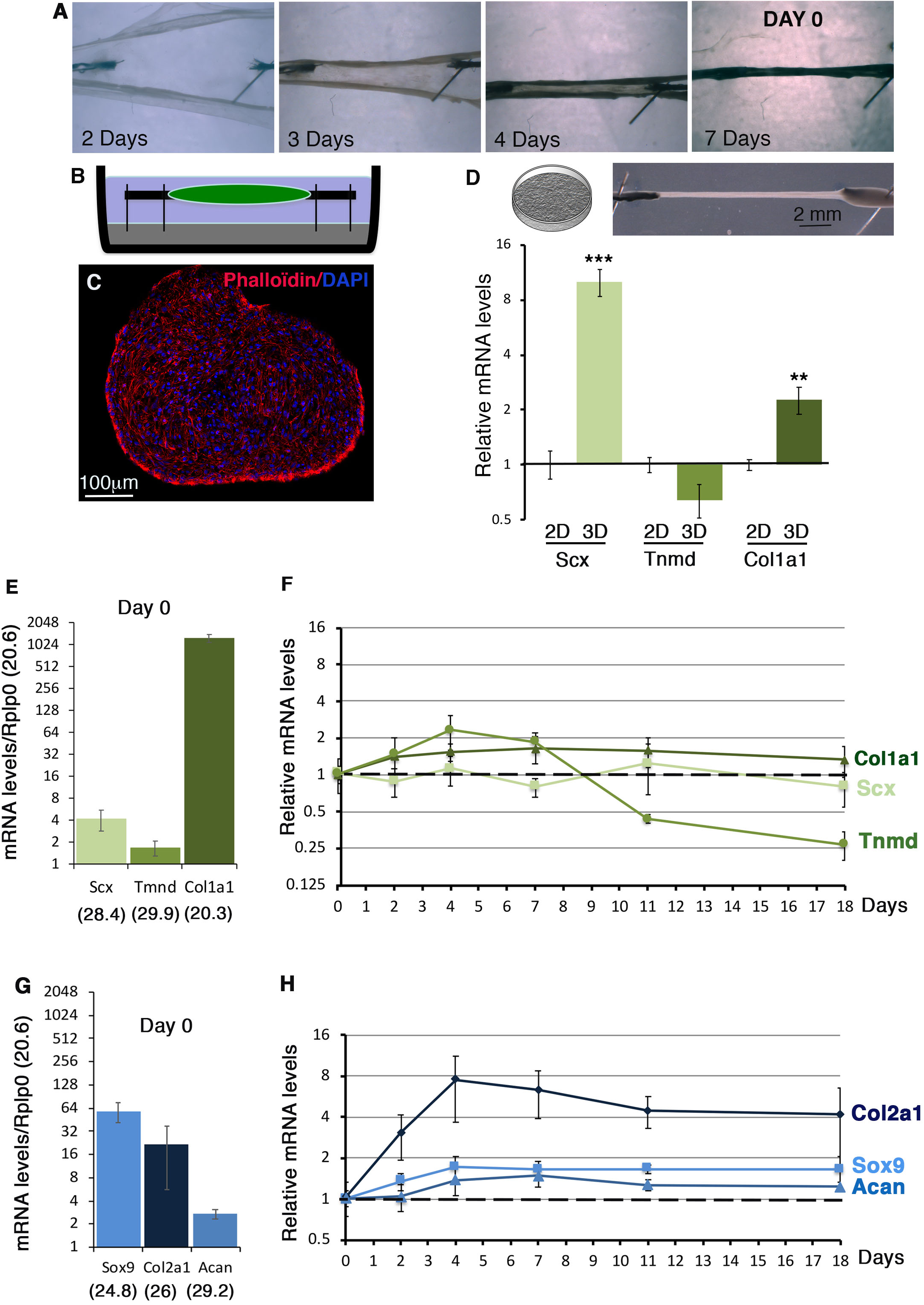
Tendon and cartilage gene expression in C3H10T1/2 cells in 3D-fibrin gel condition. **(A)** 3D-constructs were made by mixing C3H10T1/2 cells with a fibrin gel. 3 to 7 days were required to form a construct. Day 0 was considered when constructs were formed (here for 7 days). (**B**) Schematic representation of a 3D fibrin construct. (**C**) Transverse section of a construct, 24 hours after formation, labelled with DAPI and Phaloïdin. (**D**) RT-q-PCR analyses of the expression levels for the *Scx, Tnmd* and *Col1a1* tendon genes in C3H10T1/2 cells cultured in 3D-fibrin constructs compared to 2D conditions on plastic for 10 days. The relative mRNA levels were calculated using the 2^^-ΔΔCt^ method using the 2D-conditions as controls. For each gene, the mRNA levels of 2D-conditions were normalized to 1. The p-values were calculated using the Mann-Whitney test. **(E-H)** RT-q-PCR analyses of the expression levels for *Scx, Tnmd* and *Col1a1* tendon genes (E,F) and *Sox9* and *Col2a1* cartilage genes (G,H) in C3H101/2 cells cultured in 3D fibrin gel conditions over time, at Day 0 (N=4), 2 days (N=4), 4 days (N=4) 7 days (N=8), 11 days (N=4) and 18 days (N=4). (**E,G**) At Day 0, the means of the Cts (obtained from 500 ng of mRNAs) are indicated in brackets for each gene. For each gene, the ΔCts were calculated using the *Rplp0* gene as reference gene. ΔCt = Ct *gene* - Ct *Rplp0.* The mRNA levels were reported using the 2^^-ΔCt^ method. For each gene, 2^^-ΔCt^ were multiplied per 10^3^ in order to obtain values above 1. Graphs show means ± standard deviations of 4 biological samples. (**F,H**) RT-q-PCR analyses of the relative expression levels of *Scx, Col1a1* and *Tnmd* tendon genes (F) and *Sox9, Col2a1* and *Acan* cartilage genes (H) in 3D-fibrin constructs over time. The relative mRNA levels were calculated using the 2^^-ΔΔCt^ method using the Day 0 condition as controls. For each gene, the mRNA levels of Day 0 condition were normalized to 1.

Tendon and cartilage gene expression was analyzed at different time points (Day 0, Day 2, Day 4, Day 7, Day 11 and Day 18). The Day 0 time point corresponds to the day when the constructs were formed (Fig. 5A) and was the reference time point. The expression profile of tendon genes at Day 0 in 3D-fibrin constructs (Figure 5E) was similar to that in 2D-cultures (Figures 1C, 2C,D), *i.e.* relatively high levels of *Col1a1* mRNAs compared to *Scx* and *Tnmd*. In contrast to a decrease in 2D-cultures (Figures 3,4), *Scx* and *Col1a1* displayed an unchanged expression in 3D-fibrin constructs over time following that observed in Day 0 (Figure 5F). Similarly to 2D-cultures (Figure 3C, 4C), *Tnmd* expression displayed a bell shape with a maximum of 2-fold increase between Day 0 and Day 7 in 3D-fibrin constructs (Figure 5F). The cartilage genes, *Sox9* (progenitors), *Acan* and *Col2a1* (differentiated cells) were expressed at Day 0 (Fig. 5G) and increased over time in 3D-fibrin constructs, indicating that the potential of C3H10T1/2 cells to differentiate into cartilage is maintained in 3D-fibrin constructs (Fig. 5H). The expression of *Pparg* (early fat differentiation marker) was above 32 cycles at Day 7 indicating an absence of adipocyte differentiation of C3H10T1/2 cells in 3D-fibrin constructs.

We conclude that the 3D-environment in fibrin gel maintains tendon gene expression in C3H10T1/2 cells over time.

### TGFβ effect on tendon gene expression in C3H10T1/2 cells in 2D- and 3D-culture systems

The canonical TGFβ/SMAD2/3 pathway is recognized to have a pro-tenogenic effect in cell cultures based on *Scx* expression (Guerquin et al., 2013; Havis et al., 2014, 2016; Lorda-Diez et al., 2009; Pryce et al., 2009). There are not many recognized transcriptional readout of TGFβ activity, but Smad7 is a negative-feedback regulator that is considered to be a general SMAD2/3 transcriptional target gene (Massagué, 2012). We assessed the activity of TGFβ/SMAD2/3 signaling pathway with *Smad7* expression in C3H10T1/2 cells cultured with plastic and silicon substrates over time. The initial cycle threshold number for the *Smad7* gene (Ct=23.9 for plastic and Ct = 24.7 for silicone) indicated that *Smad7* was expressed in C3H10T1/2 cells, in both substrate culture conditions at Day 0. The *Smad7* expression profile displayed a mirror image to that of *Tnmd*, while *Smad7* expression followed that of *Scx* in C3H10T1/2 cells cultured in plastic and silicon substrate 2D-conditions (Figures 6A,B, 4C, 3C). The *Smad7* expression profile also mapped that of *Scx* and differed from that of *Tnmd* in 3D-fibrin constructs (Figures 6C, 5E). The expression profiles of *Tnmd* and *Scx* genes in Figures 3C, 4C, 5E was added on panels A-C of Figure 6 to facilitate comparison with Smad7 expression levels. The fact that the activity of TGFβ/SMAD2/3 signaling pathway followed that of *Scx* expression in C3H10T1/2 cells whatever the culture system was consistent with the recognized positive regulation of *Scx* expression by the TGFβ/SMAD2/3 signaling pathway in C3H10T1/2 cultures (Guerquin et al., 2013; Havis et al., 2014; Havis et al., 2016). The opposite direction of *Tnmd* and *Smad7* expression profiles in C3H10T1/2 cells (Figures 6A-C) suggested that active TGFβ/SMAD2/3 signaling pathway downregulated *Tnmd* expression.

**Figure 6.**
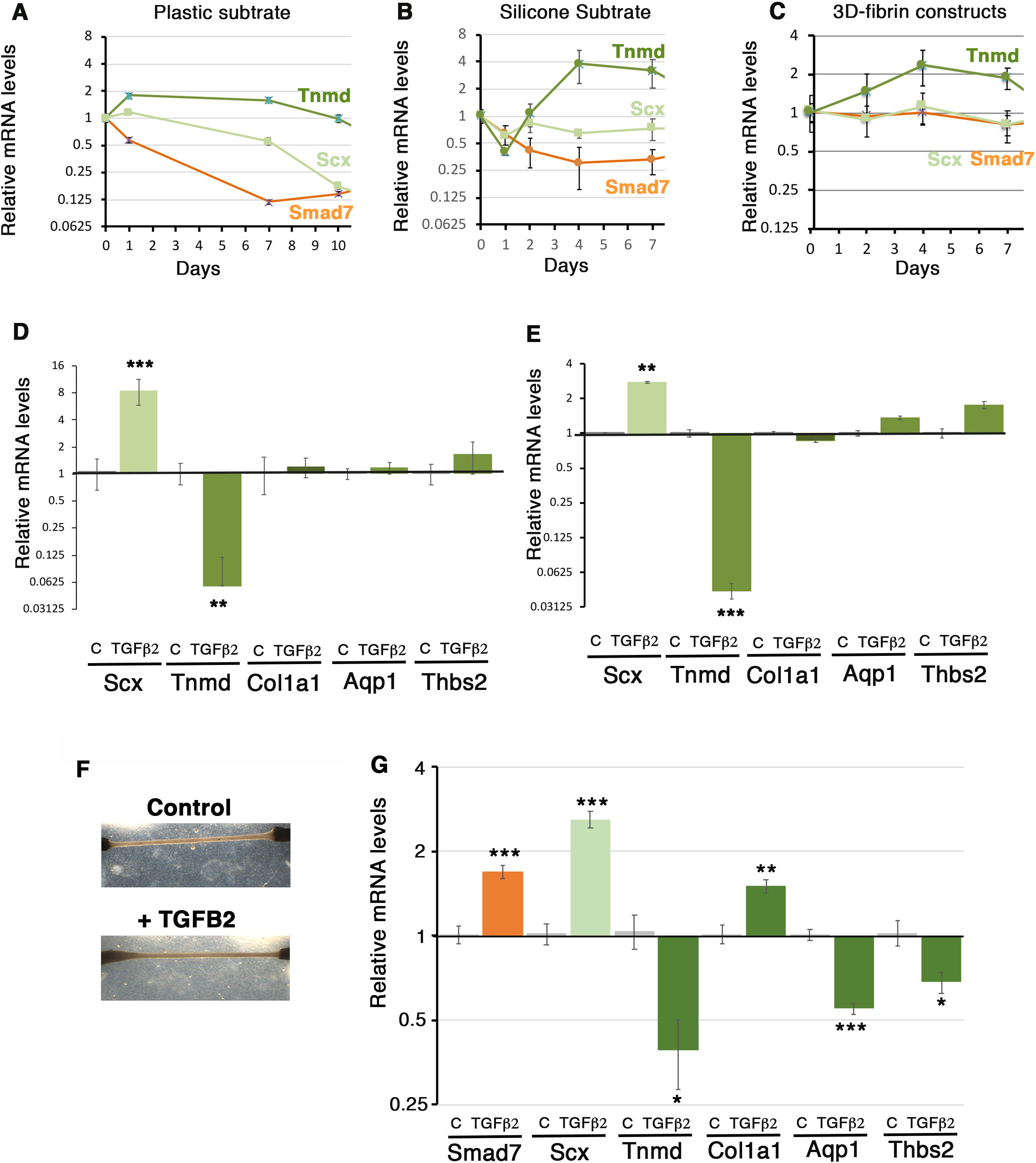
Antagonistic effect of TGFβ2 on *Tnmd* and *Scx* expression in C3H10T1/2 cells in 2D- and 3D-culture systems. (**A-C**) RT-q-PCR analyses of the expression levels of *Smad7* gene in C3H10T1/2 cells cultured upon plastic (A) and silicone (B) substrates during the expansion phase or cultured in 3D-fibrin gel condition (C). (A-C) The relative mRNA levels were calculated using the 2^^-ΔΔCt^ method. For each gene, the mRNA levels of the Day 0 condition were normalized to 1. (A) Graph shows means ± standard deviations of 4 biological samples. (B) Graph shows means ± standard deviations of 5 biological samples (C) Graph shows means ± standard deviations of 4 biological samples. The expression profiles of *Scx* and *Tnmd* shown in Figures 3C, 4C, 5F have been plotted on panels A,B,C to facilitate comparison with those of *Smad7*. (**D,E**) RT-q-PCR analyses of the expression levels of tendon gene expression in C3H10T1/2 cells cultured in control or TGFβ2-supplemented media and seeded at 10^5^ cells (11 110 cells/cm^2^) (D) and 10^4^ cells (1 110 cells/cm^2^) (E). (D,E) Graphs show mean ± standard deviations of 6 biological samples. The relative mRNA levels were calculated using the 2^^-ΔΔCt^ method. For each gene, the mRNA levels of control conditions were normalized to 1. (**F,G**) 3D-fibrin constructs in control or TGFβ2-supplemented media. (F) Images showing no significant variations in the morphology of the TGFβ2-treated 3D-constructs (below) compared to controls (above). (G) RT-qPCR analysis of the relative expression of the tendon-associated genes in 3D-constructs treated or not with TGFβ2. Graph shows mean ± standard deviations of 6 biological samples. The relative mRNA levels were calculated using the 2^^-ΔΔCt^ method. For each gene, the mRNA levels of control conditions were normalized to 1. The p-values were calculated using the Mann-Whitney test.

In order to test this, we analyzed the effect of TGFβ2 on *Tnmd* expression in C3H10T1/2 cells seeded at 2 different cell densities, 11 110 cells/cm^2^ (Figure 6D) and 1 110 cells/cm^2^ (Figure 6E). TGFβ2 treatment drastically decreased *Tnmd* mRNA levels, while increasing those of *Scx* in C3H10T1/2 cells compared to no treatment, in both seeding cell densities (Figure 6D,E). Other tendon markers such as *Col1a1, Aqp1* and *Thbs2* did not display any significant variations upon TGFβ2 exposure, indicating a differential effect of TGFβ on *Scx* and *Tnmd* expression in C3H10T1/2 cells in 2D-culture conditions. In order to test if the negative effect of TGFβ2 on *Tnmd* expression was inherent to the 2D-culture system, we also applied TGFβ2 in C3H10T1/2 cells cultured in 3D-fibrin gel. TGFβ2 was added in the culture medium of tendon constructs for 24 hours and compared to non-treated constructs harvested at the same time. No apparent differences could be observed in the morphology of the TGFβ2-treated constructs when compared to controls (Figure 6F). Consistent with the results obtained in 2D-cultures (Figure 6D,E), we found an increase in the expression of *Scx* and a concomitant decrease in *Tnmd* expression in TGFβ2-treated 3D-tendon constructs compared to control constructs (Figure 6G). *Col1a1* expression was increased as that of *Scx*, while *Aqp1* and *Thsb2* expression was decreased as that of *Tnmd* upon TGFβ2 exposure (Figure 6G). This shows that TGFβ2 has a negative effect on *Tnmd* expression, while having a positive effect on *Scx* expression in C3H10T1/2 cells cultured in 3D-culture conditions.

We conclude that TGFβ2 is a negative regulator of *Tnmd* expression in C3H10T1/2 cells in 2D- and 3D-culture systems.

## Discussion

In the present study, we analyzed the tendon differentiation potential of C3H10T1/2 cells cultured in different conditions. Our results show that C3H10T1/2 cells behave differently for tendon gene expression depending on the substrate on which they were seeded in 2D cultures and 3D-environment. We also identified TGFβ2 as a potent negative regulator of the tendon differentiation marker *Tnmd* in C3H10T1/2 cells in 2D- and 3D-culture systems.

### Tendon differentiation potential for C3HT101/2 cells cultured on plastic and silicon substrates

C3H10T1/2 cells, although they express tendon genes in 2D-cultures (Figure 1), are not preferentially committed to the tendon lineage as compared to primary tendon or ligament cells originating from native tissues. We found that the initial tendon gene profile was similar in C3H10T1/2 cells seeded on silicone and plastic substrate in non-confluent 2D conditions (Figure 2). However, the silicone substrate was more prone to maintain the tendon phenotype of C3H10T1/2 cell cultures during the expansion and post-expansion phases over time compared to plastic substrate (Figures 3,4). A way to compare substrates of different chemical composition is to look at their stiffness. The design of our study allowed us to compare two substrates, plastic (1 GPa magnitude) and silicone (5 MPa) with a 200-fold difference in stiffness on the rigid scale. The extreme rigidity of plastic substrate (1 GPa) progressively decreases the expression of *Scx*, while a relatively less rigid substrate (5 MPa) decreases *Scx* and *Col1a1* by 2-fold in 1 day but then maintains their expression over time. The silicon substrate favors the expression of the tendon differentiation marker, *Tnmd* during the expansion phase. Based on *Scx* and *Tnmd* expression, we conclude that a substrate of 5 MPa rigidity favors the tendon phenotype in C3H10T1/2 cells over time. Although the stiffness values of both substrates display a 200-fold difference, these two substrates are still in the rigid scale favorable for bone differentiation (Discher et al., 2009). Consistently, C3H10T1/2 mesenchymal stem cells cultured on these two substrates (plastic and silicone) display a significant and drastic increase in the expression of the bone differentiation marker (*bglap*) over time. Because there was no addition of bone differentiation medium in the culture conditions, we believe that cell confluence favors bone differentiation of C3H10T1/2 cells cultured on these two rigid substrates. The dramatic increase in the expression of the early differentiation fat marker *Pparg* in plastic substrate (high stiffness) is counterintuitive with the range of soft stiffness known to promote fat differentiation (Discher et al., 2009). We interpret the ability of C3H10T1/2 cells to differentiate towards the fat lineage under a stiff substrate by the fact that C3H10T1/2 cells make multilayers upon confluence. One obvious hypothesis is that cells expressing *Pparg* at 14 days of culture could be those in the superficial cell multilayer, not in contact with the plastic substrate and thus creating a soft environment.

### TGFβ is a potent negative regulator of *Tnmd* expression in C3H10T1/2 cells in 2D and 3D culture systems

Our work identifies a striking inverse correlation between *Tnmd* expression and TGFβ activity (assessed with *Smad7* expression) in C3H10T1/2 cells cultured in 2D conditions on both plastic and silicone substrates and in 3D-fibrin gel systems over time. Consistently, TGFβ2 drastically decreases *Tnmd* expression, while promoting that of *Scx* in C3H10T1/2 cells cultured in 2D- and 3D-culture systems. The opposite behavior of *Scx* and *Tnmd* expression in cell cultures over time and upon TGFβ application could reflect different steps of tenogenesis, with a progenitor step revealed by *Scx* and a differentiation one by *Tnmd*. During development, *Scx* is expressed before *Tnmd* and it has been shown that *Scx* is required and sufficient for *Tnmd* expression in developing tendons (Murchison et al., 2007; Shukunami et al., 2006). *Scx* and *Tnmd* also display opposite expression profiles in primary tendon cells over time (Shukunami et al., 2018). Moreover, Scx has been recently shown to directly regulate *Tnmd* transcription in primary tendon cells (Shukunami et al., 2018). The absence of *Tnmd* activation concomitant with *Scx* increase upon TGFβ application (Figure 6D-F) is unexpected but indicates that TGFβ inhibits *Tnmd* expression in C3H10T1/2 cells in 2D- and 3D-culture conditions. It has to be noted that TGFβ2 increased the expression of both *Scx* and *Tnmd* genes in chick and mouse limb explants (Havis et al., 2014; Havis et al., 2016), in high density cultures of chick limb cells (Lorda-Diez et al., 2009) or in 3D-culture systems made of human tendon cells (Bayer et al., 2010). We cannot exclude that the negative regulation of TGFβ on *Tnmd* expression is cell-type specific and related to mesenchymal stem cells. The relevance to the in vivo situation of *Tnmd* inhibition by TGFβ2 in C3H10T1/2 cells requires further investigation.

### Conclusion

This study shows that culture conditions such as expansion, confluence, substrates, 2D and 3D environment affect the tendon differentiation potential of a murine cell line of mesenchymal stem cells, C3H10T1/2 cells. We also identify TGFβ2 as a negative regulator of *Tnmd* expression in C3H10T1/2 cells in 2D- and 3D- culture systems. The identification of the optimum conditions that induce tendon cell differentiation *in vitro* is of particular interest in order to optimize tendon cell culture protocols from stem cells that can be used for tendon repair.

## Material and methods

### Cell cultures

The multipotent mouse mesenchymal stem cells, C3H10T1/2 cells (Reznikoff et al., 1973) were cultured on 6-wells TPP plastic culture plates (Merck) or 6-wells Uniflex Flexcell plates (FlexCell Int) made of silicone substrate coated with type I collagen, in Dulbeccos Modified Eagles Medium (DMEM, Invitrogen) supplemented with 10% fetal bovine serum (FBS, Sigma) 1% penicillin-sreptomycin (Sigma), 1% glutamin (Sigma) and incubated at 37°C in humidified atmosphere with 5% of CO_2_. The culture medium was changed every 48 hours. To study the effect of cell number on tendon gene expression, 0.5×10^5^ (5 555 cells/cm^2^), 10^5^ (11 110 cells/cm^2^) and 2×10^5^ (22 220 cells/cm^2^) C3H10T1/2 cells were seeded in 9 cm^2^ 6 wells TPP tissue culture plates (plastic substrate), left for 16 hours in culture and analyzed for tendon gene expression by RT-qPCR. 250 ng of RNA were extracted from each sample before proceeding with the RT-qPCR. For the study of the effect of the initial cell number, 6 samples (N=6) were analyzed in each condition of cell density. The *Rplp0* gene was used as the reference gene.

For the analysis of the differentiation potential of C3H10T1/2 cells seeded on plastic substrate, 10^5^ cells were seeded in 6-wells TPP culture plates (11 110 cells/cm^2^) and left for 16 hours in culture. This defined the Day 0 (N=4) and then cells were cultured for another 24 hours (1 day) (N=4), 7 days (N=4), 10 days (N=5) and 14 days (N=4). 500 ng of RNAs were extracted from each sample before proceeding with the RT-qPCR.

For the analysis of the differentiation potential of C3H10T1/2 cells seeded on silicon substrate coated with type I collagen, 10^5^ cells were seeded in 6-wells Uniflex Flexcell plates and left for 16 hours in culture. This defined the Day 0 (N=5) and then cells were cultured for another 24 hours (1 day) (N=6), 48 hours (2 days) (N=6), 7 days (N=6), 11 days (N=6). 500 ng of RNA were extracted from each sample before proceeding with the RT-qPCR.

### 3D-engineered tendon constructs in fibrin gels

3D fibrin-based tendon-like constructs made of mouse C3H10T1/2 cells were performed as previously described (Kapacee et al., 2008). Briefly, for each construct, 400 µl of cell suspension (7.5×10^5^ cells) were mixed with 20 mg/ml fibrinogen (Sigma, St Louis, MO, USA) and 200 U/ml thrombin (Sigma, St Louis, MO, USA). The fibrin gels containing cells were seeded in prepared SYLGARD-covered wells (DowChemical, Midland, MI, USA), in which two 8 mm- sutures (Ethican, Sommerville, NJ, USA) were pinned 10 mm apart. Culture medium containing 200 µM of L-ascorbic acid 2-phosphate was added to the wells and gels were scored every day for a proper contraction into a linear construct. After 5 to 7 days, the C3H10T1/2 cells formed continuous tendon-like constructs between the 2 anchors. This was considered as Day 0. Each tendon construct was considered as a biological sample. The mRNA levels of each construct were analyzed by q-RT-PCR at 2 days, 4 days, 7 days, 11 days and 18 days after Day 0.

### TGF-β treatment on 2D and 3D cultures

10^5^ or 10^4^ C3H10T1/2 cells were plated on 6-wells TPP culture plates (plastic) and grown for 40 hours. Then, human recombinant TGFβ2 (RD System) was applied at 20 ng/ml to C3H10T1/2 cells for 24 hours. Cells were grown for another 24 hours without TGFβ2 supplementation in the medium. Control cells were treated with Bovin Serum Albumin and HCl (BSA-HCl) in the same volume than that applied for TGFβ2 treatment. TGF-β2-treated and non-treated C3H10T1/2 cells were then fixed and processed for q-PCR assays to analyze gene expression. In each condition, 4 biological samples (N=4) were used.

3D tendon constructs were treated with TGFβ2 or with BSA-HCl (controls) at Day 7 of culture, for 24 hours. In each condition, 5 biological samples (N=5) were used for q-PCR analysis.

### RNA isolation, Reverse transcription and Real Time quantitative-PCR

Total RNAs were extracted from 2D and 3D cell cultures: C3H10T1/2 cells cultured on classic culture dishes at Day 0, 1 day, 7 days, 10 days and 14 days, C3H10T1/2 cells cultured on silicone substrate at Day 0, 1 day, 2 days, 4 days, 7 days and 11 days, C3H10T1/2 cells cultured in 3D fibrin gel conditions at Day 0, 2 days, 4 days, 7 days, 11 days and 18 days, and TGFβ2-treated C3H10T1/2 cells cultured in 2D and 3D conditions. Total RNA was isolated using the RNeasy mini kit (Qiagen) with 15 min of DNase I (Qiagen) treatment according to the manufacturer’s protocol. For RT-qPCR analyses, 250 ng or 500 ng RNAs was Reverse-Transcribed using the High Capacity Retrotranscription kit (Applied Biosystems). RT-qPCR PCR was performed using SYBR Green PCR Master Mix (Applied Biosystems) using primers listed in Table 1. We used as housekeeping genes, the *rn18s* (other named 18S) and *Rplp0* (other named 36b4) genes as housekeeping genes. The *rn18s* and *Rplp0* genes did not show any variation in the different experimental conditions. The *Rplp0* gene is detected around a Ct (threshold cycle) of 19.5 for 250 ng of RNAs and around a Ct of 18.5 Ct for 500 ng of RNAs. This result is consistent with the log2-linear plot of the PCR signal. A decrease of one cycle corresponds to a fold-2 increase of RNAs (Livak and Schmittgen, 2001). The *rn18s* gene was detected around 7.5 Ct for 500 ng of RNA. The relative mRNA levels were calculated using the 2^-ΔΔCt method (Livak and Schmittgen, 2001; Schmittgen and Livak, 2008). The ΔCt values were obtained by calculating the differences: Ct(gene of interest) – Ct(housekeeping gene) in each sample. ΔΔCt values were obtained by calculating the differences between ΔCt (Experimental condition) and the average of control ΔCt values. For the analysis of the relative mRNA levels of cells cultured over time in classic culture plates (plastic substrate), Uniflex Flexcell plates (silicone substrate) or 3D-fibrin condition, the values of the Day 0 time points were considered as controls and were normalized to 1. For the relative mRNA level analysis in TGFβ2-treated cells in 2D or 3D conditions, the cells in the absence of TGFβ2 supplementation were considered as controls and were normalized to 1. For the absolute quantification of gene expression, 16 hours after plating 10^5^ cells, Y-axes correspond to 2^-ΔCtx10^3^ against the *Rplp0* house keeping gene from 250 ng of RNA (Figure 1C), to 2^-ΔCtx10^3^ against the *Rn18S* house keeping gene from 500 ng of RNA (Figure 2C) and 2^-ΔCtx10^4^ against the *Rplp0* house keeping gene from 500 ng of RNA (Figure 2D).

**Table 1:**
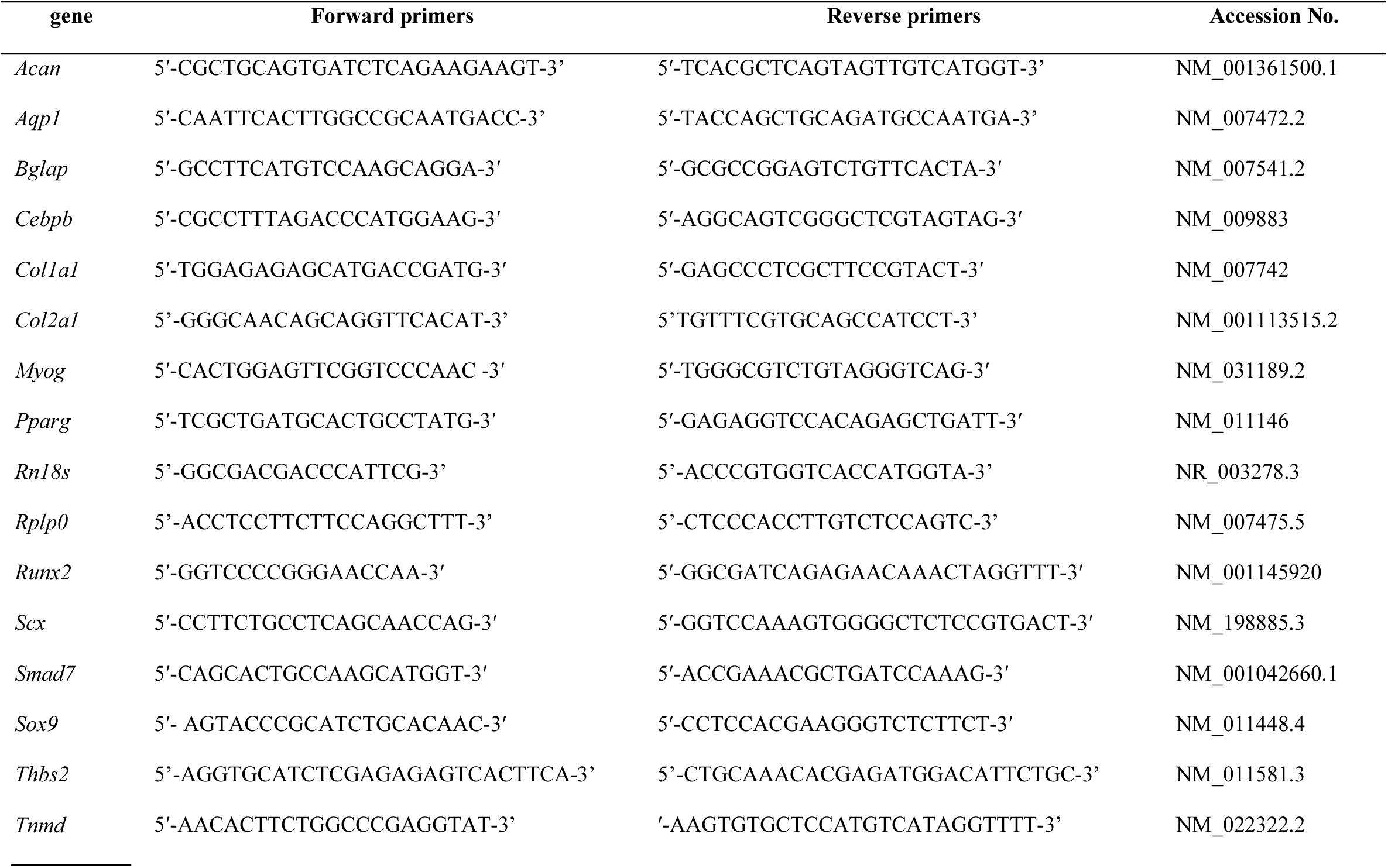
Primers used for Real-time quantitative PCR.

### Statistical analyses

Results are shown as means ± standard deviation (SD). The exact number of independent biological samples (4 to 6) is reported for each experiment. RT-q-PCR data were analyzed with the non-parametric Mann-Withney test with Graphpad Prism V6. The asterisks in histograms indicate *p* values that was considered significant, *<0.05, **<0.01, ***<0.001.

## Acknowledgements

We thank Sophie Gournet for illustrations. We are grateful to lab members for reading the manuscript.

## Author contributions

This study was conceived by DD and supervised by DD and MM. Experiments were performed by LG, MAB, IC, MO. Data analysis and interpretation was performed by LG, MM and DD. DD and MM acquired funding. The manuscript was written by LG, MM and DD with involvement and final approval by all co-authors

## Bibliography

Abo-Aziza, F. A. M. and A A, Z. (2017). The Impact of Confluence on Bone Marrow Mesenchymal Stem (BMMSC) Proliferation and Osteogenic Differentiation. Int. J. Hematol. stem cell Res. 11, 121–132.

Alberton, P., Dex, S., Popov, C., Shukunami, C., Schieker, M. and Docheva, D. (2015). Loss of Tenomodulin Results in Reduced Self-Renewal and Augmented Senescence of Tendon Stem/Progenitor Cells. Stem Cells Dev. 24, 597–609.

Bayer, M. L., Yeung, C.-Y. C., Kadler, K. E., Qvortrup, K., Baar, K., Svensson, R. B., Peter Magnusson, S., Krogsgaard, M., Koch, M. and Kjaer, M. (2010). The initiation of embryonic-like collagen fibrillogenesis by adult human tendon fibroblasts when cultured under tension. Biomaterials 31, 4889–4897.

Bellas, E. and Chen, C. S. (2014). Forms, forces, and stem cell fate. Curr. Opin. Cell Biol. 31, 92–7.

Buckingham, M. (2017). Gene regulatory networks and cell lineages that underlie the formation of skeletal muscle. Proc. Natl. Acad. Sci. 114, 5830–5837.

Caplan, A. I. (1991). Mesenchymal stem cells. J. Orthop. Res. 9, 641–650.

Dex, S., Lin, D., Shukunami, C. and Docheva, D. (2016). TENOgenic MODULating INsider factor: systematic assessment on the functions of tenomodulin gene. Gene 587, 1–17.

Dex, S., Alberton, P., Willkomm, L., Söllradl, T., Bago, S., Milz, S., Shakibaei, M., Ignatius, A., Bloch, W., Clausen-Schaumann, H., et al. (2017). Tenomodulin is Required for Tendon Endurance Running and Collagen I Fibril Adaptation to Mechanical Load. EBioMedicine 20,.

Discher, D. E., Mooney, D. J. and Zandstra, P. W. (2009). Growth Factors, Matrices, and Forces Combine and Control Stem Cells. Science (80-.). 324, 1673–1677.

Docheva, D., Hunziker, E. B., Fässler, R. and Brandau, O. (2005). Tenomodulin is necessary for tenocyte proliferation and tendon maturation. Mol. Cell. Biol. 25, 699–705.

Engler, A. J., Sen, S., Sweeney, H. L. and Discher, D. E. (2006). Matrix Elasticity Directs Stem Cell Lineage Specification. Cell 126, 677–689.

Gaut, L. and Duprez, D. (2016). Tendon development and diseases. Wiley Interdiscip. Rev. Dev. Biol. 5, 5–23.

Gaut, L., Robert, N., Delalande, A., Bonnin, M.-A., Pichon, C. and Duprez, D. (2016). EGR1 Regulates Transcription Downstream of Mechanical Signals during Tendon Formation and Healing. PLoS One 11, e0166237.

Guerquin, M. J., Charvet, B., Nourissat, G., Havis, E., Ronsin, O., Bonnin, M. A., Ruggiu, M., Olivera-Martinez, I., Robert, N., Lu, Y., et al. (2013). Transcription factor EGR1 directs tendon differentiation and promotes tendon repair. J. Clin. Invest. 123, 3564–3576.

Havis, E., Bonnin, M.-A., Olivera-Martinez, I., Nazaret, N., Ruggiu, M., Weibel, J., Durand, C., Guerquin, M.-J., Bonod-Bidaud, C., Ruggiero, F., et al. (2014). Transcriptomic analysis of mouse limb tendon cells during development. Development 141, 3683–96.

Havis, E., Bonnin, M. M.-A., De Lima, J. J. E., Charvet, B., Milet, C. and Duprez, D. (2016). TGFβ and FGF promote tendon progenitor fate and act downstream of muscle contraction to regulate tendon differentiation during chick limb development. Development 143, 3839–3851.

Hsieh, C.-F., Yan, Z., Schumann, R., Milz, S., Pfeifer, C., Schieker, M. and Docheva, D. (2018). In Vitro Comparison of 2D-Cell Culture and 3D-Cell Sheets of Scleraxis-Programmed Bone Marrow Derived Mesenchymal Stem Cells to Primary Tendon Stem/Progenitor Cells for Tendon Repair. Int. J. Mol. Sci. 19, 2272.

Huang, A. H., Lu, H. H. and Schweitzer, R. (2015). Molecular regulation of tendon cell fate during development. In Journal of Orthopaedic Research, pp. 800–812.

Humphrey, J. D., Dufresne, E. R. and Schwartz, M. A. (2014). Mechanotransduction and extracellular matrix homeostasis. Nat. Rev. Mol. Cell Biol. 15, 802–812.

Ivanovska, I. L., Shin, J.-W., Swift, J. and Discher, D. E. (2015). Stem cell mechanobiology: diverse lessons from bone marrow. Trends Cell Biol. 25, 523–532.

Kapacee, Z., Richardson, S. H., Lu, Y., Starborg, T., Holmes, D. F., Baar, K. and Kadler, K. E. (2008). Tension is required for fibripositor formation. Matrix Biol. 27, 371–375.

Kapacee, Z., Yeung, C.-Y. C., Lu, Y., Crabtree, D., Holmes, D. F. and Kadler, K. E. (2010). Synthesis of embryonic tendon-like tissue by human marrow stromal/mesenchymal stem cells requires a three-dimensional environment and transforming growth factor β3. Matrix Biol. 29, 668–677.

Karsenty, G., Kronenberg, H. M. and Settembre, C. (2009). Genetic Control of Bone Formation. Annu. Rev. Cell Dev. Biol. 25, 629–648.

Kilian, K. A., Bugarija, B., Lahn, B. T. and Mrksich, M. (2010). Geometric cues for directing the differentiation of mesenchymal stem cells. Proc. Natl. Acad. Sci. U. S. A. 107, 4872–7.

Kuo, C. K., Marturano, J. E. and Tuan, R. S. (2010). Novel strategies in tendon and ligament tissue engineering: Advanced biomaterials and regeneration motifs. BMC Sports Sci. Med. Rehabil. 2, 20.

Liu, H., Zhang, C., Zhu, S., Lu, P., Zhu, T., Gong, X., Zhang, Z., Hu, J., Yin, Z., Heng, B. C., et al. (2015). Mohawk Promotes the Tenogenesis of Mesenchymal Stem Cells Through Activation of the TGFβ Signaling Pathway. Stem Cells 33, 443–455.

Liu, C.-F., Samsa, W. E., Zhou, G. and Lefebvre, V. (2017). Transcriptional control of chondrocyte specification and differentiation. Semin. Cell Dev. Biol. 62, 34–49.

Livak, K. J. and Schmittgen, T. D. (2001). Analysis of Relative Gene Expression Data Using Real-Time Quantitative PCR and the 2−ΔΔCT Method. Methods 25, 402–408.

Lorda-Diez, C. I., Montero, J. A., Martinez-Cue, C., Garcia-Porrero, J. A. and Hurle, J. M. (2009). Transforming Growth Factors β Coordinate Cartilage and Tendon Differentiation in the Developing Limb Mesenchyme. J. Biol. Chem. 284, 29988–29996.

Maeda, T., Sakabe, T., Sunaga, A., Sakai, K., Rivera, A. L., Keene, D. R., Sasaki, T., Stavnezer, E., Iannotti, J., Schweitzer, R., et al. (2011). Conversion of Mechanical Force into TGF-β-Mediated Biochemical Signals. Curr. Biol. 21, 933–941.

Mammoto, T., Mammoto, A. and Ingber, D. E. (2013). Mechanobiology and Developmental Control. Annu. Rev. Cell Dev. Biol. 29, 27–61.

Marturano, J. E., Schiele, N. R., Schiller, Z. A., Galassi, T. V., Stoppato, M. and Kuo, C. K. (2016). Embryonically inspired scaffolds regulate tenogenically differentiating cells. J. Biomech. 49, 3281–3288.

Massagué, J. (2012). TGFβ signalling in context. Nat. Rev. Mol. Cell Biol. 13, 616–630.

Mendias, C. L., Gumucio, J. P., Bakhurin, K. I., Lynch, E. B. and Brooks, S. V. (2012). Physiological loading of tendons induces scleraxis expression in epitenon fibroblasts. J. Orthop. Res. 30, 606–612.

Murchison, N. D., Price, B. a, Conner, D. a, Keene, D. R., Olson, E. N., Tabin, C. J. and Schweitzer, R. (2007). Regulation of tendon differentiation by scleraxis distinguishes force-transmitting tendons from muscle-anchoring tendons. Development 134, 2697–2708.

Nourissat, G., Berenbaum, F. and Duprez, D. (2015). Tendon injury: from biology to tendon repair. Nat. Rev. Rheumatol. 11, 223–233.

Pittenger, M. F., Mackay, A. M., Beck, S. C., Jaiswal, R. K., Douglas, R., Mosca, J. D., Moorman, M. A., Simonetti, D. W., Craig, S. and Marshak, D. R. (1999). Multilineage Potential of Adult Human Mesenchymal Stem Cells. Science (80-.). 284, 143–147.

Prockop, D. J. (1997). Marrow Stromal Cells as Stem Cells for Nonhematopoietic Tissues. Science (80-.). 276, 71–74.

Pryce, B. A., Watson, S. S., Murchison, N. D., Staverosky, J. A., Dunker, N. and Schweitzer, R. (2009). Recruitment and maintenance of tendon progenitors by TGF signaling are essential for tendon formation. Development 136, 1351–1361.

Ren, J., Wang, H., Tran, K., Civini, S., Jin, P., Castiello, L., Feng, J., Kuznetsov, S. A., Robey, P. G., Sabatino, M., et al. (2015). Human bone marrow stromal cell confluence: effects on cell characteristics and methods of assessment. Cytotherapy 17, 897–911.

Reznikoff, C. a, Brankow, D. W. and Heidelberger, C. (1973). Establishment and Characterization of a Cloned Line of C3H Mouse Embryo Cells Sensitive to Postconfluence Inhibition of Division Establishment and Characterization of a Cloned Line of C3H Mouse Embryo Cells Sensitive to Postconfluence Inhibition of. Cancer Res. 3231–3238.

Schiele, N. R., Marturano, J. E. and Kuo, C. K. (2013). Mechanical factors in embryonic tendon development: potential cues for stem cell tenogenesis. Curr. Opin. Biotechnol. 24, 834–840.

Schmittgen, T. D. and Livak, K. J. (2008). Analyzing real-time PCR data by the comparative C(T) method. Nat. Protoc. 3, 1101–8.

Schweitzer, R., Chyung, J. H., Murtaugh, L. C., Brent, A. E., Rosen, V., Olson, E. N., Lassar, A. and Tabin, C. J. (2001). Analysis of the tendon cell fate using Scleraxis, a specific marker for tendons and ligaments. Development 128, 3855–3866.

Schweitzer, R., Zelzer, E. and Volk, T. (2010). Connecting muscles to tendons: tendons and musculoskeletal development in flies and vertebrates. Development 137, 2807–2817.

Shukunami, C., Takimoto, A., Oro, M. and Hiraki, Y. (2006). Scleraxis positively regulates the expression of tenomodulin, a differentiation marker of tenocytes. Dev. Biol. 298, 234–247.

Shukunami, C., Takimoto, A., Nishizaki, Y., Yoshimoto, Y., Tanaka, S., Miura, S., Watanabe, H., Sakuma, T., Yamamoto, T., Kondoh, G., et al. (2018). Scleraxis is a transcriptional activator that regulates the expression of Tenomodulin, a marker of mature tenocytes and ligamentocytes. Sci. Rep. 8, 3155.

Yao, L., Bestwick, C. S., Bestwick, L. A., Maffulli, N. and Aspden, R. M. (2006). Phenotypic Drift in Human Tenocyte Culture. Tissue Eng. 12, 1843–1849.

Yeung, C.-Y. C., Zeef, L. A. H., Lallyett, C., Lu, Y., Canty-Laird, E. G. and Kadler, K. E. (2015). Chick tendon fibroblast transcriptome and shape depend on whether the cell has made its own collagen matrix. Sci. Rep. 5, 13555.

Yoshimoto, Y., Takimoto, A., Watanabe, H., Hiraki, Y., Kondoh, G. and Shukunami, C. (2017). Scleraxis is required for maturation of tissue domains for proper integration of the musculoskeletal system. Sci. Rep. 7, 45010.

Zhang, Y.-J., Chen, X., Li, G., Chan, K.-M., Heng, B. C., Yin, Z. and Ouyang, H.-W. (2018). Concise Review: Stem Cell Fate Guided By Bioactive Molecules for Tendon Regeneration. Stem Cells Transl. Med. 7, 404–414.

